# Effect of the BMPR-II-SMAD3/MRTF complex on proliferation and migration of ASMCs and the mechanism in asthma

**DOI:** 10.1101/2020.11.19.389585

**Authors:** Wenbo Gu, Jiahui Lei, Jing Xie, Yali Xiao, Zhenping Zhang, Limin Zhao

## Abstract

Phenotype transformation of airway smooth muscle (ASM) is the key feature of airway remodelling in asthma. A variety of smooth muscle-specific genes and proteins, including SMAD3, BMPR-II, and MRTF, are involved in airway remodelling in asthma. Bone morphogenetic protein (BMP) signalling plays an important role in the physiological and pathological processes of asthma. As a receptor of BMP signalling, BMPR-II has important roles in variety of cellular processes. However, the understanding of the roles and underlying mechanism of BMPR-II in airway smooth muscle cells (ASMCs) in asthma remains incomplete. First, we observed significant increases in BMPR-II, SMAD3 and MRTF in asthmatic ASMCs at both the mRNA and protein levels. Second, we observed that silencing of siBMPR-II and siMRTF inhibits proliferation, migratory capacity and intracellular Ca^2+^ concentration in ASMCs. Furthermore, our results revealed these three factors, BMPR-II, SMAD3 and MRTF, form a complex that affects the bioactivities of ASMCs. Taken together, this study indicates that the BMPR-II-SMAD3/MRTF signalling pathway is involved in the process of ASM remodelling, providing novel avenues for the identification of new therapeutic modalities.

## Introduction

Asthma is a common chronic respiratory disease that poses a serious potential threat to patients’ physical and mental health [1]. Meanwhile, it also causes significant financial burdens to patients and their families [2]. Symptoms of asthma attacks primarily include chest tightness, wheezing, short of breath, and coughing. These symptoms are closely correlated with variable airflow limitation, airway inflammation, airway hyperresponsiveness, and airway remodelling [2]. Previous studies have claimed that airway inflammation is a prerequisite of airway remodelling [3]. However, this view has been recently debated by some scholars who posit that airway remodelling may occur prior to airway inflammation and may be the leading cause of asthma attacks [3]. Studies have shown that overexpression of ORMDL3 may induce abnormal proliferation and contractility of ASMCs in the absence of airway inflammation [4]. This research further confirmed that abnormal expression of specific gene and proteins in ASMCs may be the primary event in asthma rather than a secondary response to airway inflammation [4]. Clonal findings have verified that anti-inflammatory therapy, while improving the symptoms and controlling acute episodes, does not change the nature of airway remodelling. Therefore, many patients experience refractory asthma [5]. The results above all confirmed that airway remodelling plays a pivotal role in the pathogenesis of asthma. To overcome this disease, it is essential to identify an effective therapy to reverse airway remodelling.

Airway remodelling refers to changes in structure and function of the airway wall [6]. Pathological airway remodelling primarily includes epithelium shedding, subepithelial fibrosis, goblet cell hyperplasia, and ASMC hyperplasia and hypertrophy [6]. Recently, increasing research has found that the mass of the ASM layer in asthmatic airway tissue is significantly thicker than that in non-asthmatics [7]. Biopsies from patients with severe asthma reveal larger ASM areas and shorter distance between the subepithelial basement membrane and airway smooth muscle. Both ASM zone and SBM-ASM distance are closely related to FEV1 before and after use of bronchodilators in asthma patients, indicating that ASM content may be closely related to the severity of airway obstruction and may represent one of the critical determinants of disease severity [8]. The increase in ASM mass is classically defined as increased cell numbers resulting from augmented proliferation and decreased apoptosis, as well as cellular hypertrophy. Meanwhile, ASMCs become hypertrophic and other cells differentiate into ASMCs and migrate to into the ASM bundle, which also contributes significantly to augmented ASM content [9]. Nevertheless, the specific mechanisms mediating the bioactivity of ASMCs remain poorly understood.

Recently, several reports in the literature have correlated certain signalling molecules, include SMAD3, BMPR-II, and MRTF, to asthma progression. SMAD3 dependent TGF-β1 signalling has been validated to play a predominant role in airway remodelling in asthma [10]. Wang Meng et al discovered that Rhynchophylline (RHY) inhibits aberrant ASMC proliferation and alleviates airway inflammation, as well as allergic symptoms, by blocking TGF-β1-mediated Smads and MAPK signalling [10]. Previous studies indicated that BMP signalling plays a critical role in normal lung development [11]. BMP ligand signals are transmitted downstream by phosphorylating R-Smads, including Smad2 and Smad3, and are involved in modulating gene expression in the nucleus [12, 13]. M. Lynn et al’s research showed lower expression of BMPR-II on airway epithelium in mild asthma [14]. However, expression patterns on ASMCs in asthma patients remain unclear.

Several reports have previously demonstrated that TGF-β1 promotes MRTF nuclear translocation through a WNT-11 dependent mechanism and accelerates expression of smooth-α-actin through the Rho-actin-MRTF axis, further aggravating ASM remodelling [15]. During the progress of lens epithelial-mesenchymal transformation (EMT), Smad3 inhibitor treatment restrained MRTF-A nuclear translocation induced by TGF-β and reduced expression of α-SMA, revealing that Smad3 is an upstream regulator of MRTF-A nuclear translocation and that the two together regulate expression of α-SMA [16]. However, Previous study found that loss of functional BMPR-II induces loss of SMAD3, which directly promotes the proliferation and migration of pulmonary arterial smooth muscle cells (PASMCs) and indirectly induces hypertrophy of PASMCs by augmenting α-SMA expression via MRTF-dependent signalling [17]. Taken together, we concluded that the interactions among BMPR-II, SMAD3, and MRTF participate in regulating the bioactivities of various cell types. Moreover, our preliminary experiments demonstrated high expression of BMPR-II, SMAD3, and MRTF in ASMCs. Nevertheless, it remains unclear how these three factors interact and whether they are involved in regulating the bioactivities of ASMCs.

The aim of the study was to investigate the functional interaction among BMPR-II, SMAD3, and MRTF in ovalbumin-induced asthmatic rats, and to explore whether an upstream and downstream relationship exists among these three factors in primary ASMCs.

## Materials and methods

### Preparation of asthma model

All animal experiments were conducted in accordance with the International Animal Ethics Committee and the Animal Ethics Guidelines of Henan University of Traditional Chinese Medicine. Twenty SPF Wistar rats (body weight 200 ± 20 g, half male and female) were provided by the Animal Experiment Center of Henan University of Traditional Chinese Medicine. Rats were kept in sterile cages at a constant temperature of 20-25°C and a relative humidity of 50-70%. All rats were provided filtered water and pellet feed *ad libitum*. Twenty rats were randomly assigned to either the control or asthma group with 10 rats per group.

To create an asthma model, rats were intraperitoneally injected with 100 μg ovalbumin antigen suspension (Sigma-Aldrich, USA) and 1 mg aluminium hydroxide (Sigma-Aldrich, USA) on day 1, 7, and 15. Starting on the 21st day of the experiment, rats were placed in a closed container and stimulated with 5% OVA nebulized inhalation, for 30 minutes each, 1 ml/min, once/day for 6 weeks. The normal control group was intraperitoneally injected with the same amount of normal saline on the same schedule as the asthma group, and the same amount of normal saline was used for atomization.

### Isolation of primary ASMCs

Within 24 hours after the end of the last nebulization challenge, rats were anesthetized and sacrificed by intraperitoneal injection of 20% urethane at a dose of 5 ml/kg. Part of the trachea was obtained and fixed in 4% paraformaldehyde for haematoxylin-eosin staining (HE) and analysis (ServiceBio, Wuhan, China), and histological images were used to analyse pathophysiological changes in tracheal tissue structure in asthmatic rats under 200 × magnification.

The remaining tissue samples were used for isolation of primary ASMCs. Briefly, after removing the connective tissue and fat around the trachea under a microscope, tissues were cut into 1 mm×1 mm×1 mm pieces, digested with collagenase I at 37° for 1 h, and centrifuged at 200 rpm for 5 min. Then, pellets were placed in complete DMEM (Junxin biotech, Suzhou, China) containing 10% foetal bovine serum (FBS, Gibco, USA), 100 unit/ml penicillin and 100 mg/ml streptomycin (FBS, Gibco, USA) in an incubator containing 5% CO2 at 37°C. Media were changed every 3 days until the cell confluence reached 90%, and then cells were passaged using 0.25% trypsin-EDTA solution and used for experiments within 3-8 generations.

### Immunofluorescence assay

After their respective treatments, cells were washed with PBS three times and fixed in pre-cooled 4% paraformaldehyde (PFA) at room temperature (RT) for 30 min and permeabilised with 0.05% Triton at RT for 5 min. Subsequently, cells were blocked in 1% BSA for 1 h at RT. Then, cells were incubated in anti-α-SMA antibody (Abcam, Cambridge, UK) overnight at 4° followed by incubation with goat anti-rabbit antibody (Abcam, USA) at RT for 1 h. After staining with Hoechst (Junxin biotech, Suzhou, China) at 37° for 5-10 min, coverslips fluorescent sealant were applied to slides, which were then examined using a microscope (Olympus, Japan).

### Cell proliferation assays

Cell proliferation was measured using the Cell Counting Kit-8 assay (CCK8, Junxin, Suzhou, China) according to the manufacturer’s instructions. Cells were plated into 96-well plates at a concentration of 0.2×10^5^ cells/well. After their respective treatments, 10 μL CCK8 agents were added into each well, and cells were incubated for 2 hr followed by measurement of the absorbance 450 nm using a microplate reader (Thermo).

### Transwell assays

Cell migration ability was tested in Transwell chambers (Corning, USA). ASMCs were digested using 0.25% trypsin, 400 μL serum culture medium was added to Transwell chambers of a 24-well plate, and cells at a concentration of 2 × 10^5^/ml in serum-free medium were added to Transwell chamber inner chambers in 100 μL. Next, 600 μl DMEM/F-12 media containing 10% FBS was placed into each outer chamber as a chemoattractant. After incubation at 37°C for 24 h, non-migrated cells on the upper surface of the filter were removed with a cotton swab. After cells that were adhered to the lower surface of the filter were fixed in paraformaldehyde, 0.1% crystal violet stain was applied for 20 minutes. Finally, the number of stained cells was counted under a microscope.

### RNA extraction and quantitative RT-PCR

Total RNA was extracted from ASMCs using an RNA extraction kit (Junxin, Suzhou, China). cDNA was isolated according to the instructions of the cDNA synthesis kit (Invitrogen, Carlsbad, CA). Real-time quantitative PCR (qPCR) was performed using SYBR Select Master Mix (Junxin, Suzhou, China). PCR reaction conditions were as follows: 95°C pre-denaturation for 10 min, followed by 40 denaturation cycles, 95°C denaturation for 10 seconds, and 60°C denaturation for 30 seconds. PCR amplification products were analysed on an ABI 7900 fast thermal cycler (Applied Biosystems, ABI). β-actin was measured as a control. Data were calculated using the 2^–ΔΔCt^ method. Primers used in this study were synthesized by Sangon Biotech (Shanghai, China) and listed below: BMPR-II (forward primer): 5’-TCCGGGCAGGATAAATCAGGA-3’, and BMPR-II (reverse primer); 5’-GATTCTGGGAAGCAGCCGTA-3’. SMAD3 (forward primer): 5’-CGCATGAGCTTCGTCAAAGG-3’, and SMAD3 (reverse primer); 5’-CCGATCCCTTTACTCCCAGTG-3’. MRTF (forward primer): 5’-CTTTCTCAGCTCCCAATGGCT-3’, and MRTF (reverse primer); 5’-ACTTCTCGCTCGCAGACTTC-3’. β-actin (forward primer): 5’-ATGGATGACGATATCGCTGC-3’,and and β-actin (reverse primer);5’-CTTCTGACCCATACCCACCA-3’.

### Western blot analysis

ASMC protein was isolated using RIPA buffer (Sangon biotech, Shanghai, China), and the protein concentration was determined using the BCA method (Yeasen Biotech, Shanghai, China). Thirty micrograms of protein per well were run on 10% SDS-polyacrylamide (sodium dodecyl sulphate polyacrylamide) gel electrophoresis and then transferred to PVDF membranes (Millipore, USA). Blots were blocked in 5% bovine serum albumin at room temperature for 1 hour and incubated with primary antibodies overnight at 4° followed by incubation with corresponding secondary antibodies at room temperature for 1 h. Then, membranes were exposed to ECL reagent (Junxin, Suzhou, China). β-actin was measured as a loading control. Antibodies used in this study are listed below: anti-BMPR-II antibody (Abcam 130206), anti-SMAD3 antibody (Abcam 208182), anti-pSMAD3 antibody (Abcam 118825), anti-MRTF antibody (Abcam 49311) and anti-β-actin antibody (Abcam 8226) was obtained from Abcam (Cambridge, UK) and operated according to the manufacturer’s instructions.

### Cell transfection

Small interfering RNA (siRNA) was designed based on the known cDNA sequences of BMPR-II and MRTF D in the rat gene library and synthesized by GenePharma (Shanghai, China). The sequence were listed below: siBMPR-II-1: 5’-GACGCAUGGAGUAUUUGCUUGUGAU-3’; siBMPR-II-2: 5’-CAGCUGACAGAAGAAGACUUGGAAA-3’; siBMPR-II-3: 5’-CAGUCCUGAUGAACAUGAACCUUUA-3’; siMRTF-1: 5’-GCACAUGGAUGAUCUGUUUTT-3’; siMRTF-2: 5’-GCCUCCGUUAACACCACAATT-3’; siMRTF-3: 5’-GGUAUUUAUUCAAAGUCCAUCAAAU-3’; siNC: 5’-UUCUCCGAACGUGUCACGUTT-3’. SMAD3 overexpression vectors were constructed in Suzhou Junxin Biotechnology Co., Ltd (Suzhou, China).

siBMPR-II, siMRTF, siNC, pcDNA-SMAD3 and pcDNA-NC were transfected into ASMCs using Lipofectamine 3000 (Invitrogen, USA) according to the manufacturer’s protocol. About 48 hr after transfection, cells were harvested for further analysis.

### Detection of intracellular Ca2+ fluorescence by laser confocal microscopy

Intracellular Ca2+ concentrations were detected according to the instructions of the green fluorescent indicator Fluo-4AM (Yeasen Biotech, Shanghai, China) protocol. Briefly, ASMCs were plated on coverslips and used for detection after 48 hours of growth. Before detection, cells were treated with 4 μM Fluo4-AM (Sigma, Japan) in serum-free media and incubated at 37°C for 30 min. The blank group was treated with addition of a corresponding volume of D-Hank’s solution. Results were observed under an Olympus IX-70 laser confocal microscope (Olympus, Japan).

### Co-Immunoprecipitation (Co-IP)

Co-IP assays were performed using a Co-Immunoprecipitation kit (Thermofisher, USA) according to the manufacturer’s instructions. Briefly, cells were rinsed twice in ice-cold PBS and then placed in 1 ml pre-chilled lysis buffer (20 mM Tris PH 7.5, 150 mM NaCl, 1 mM EDTA, 1% NP-40, 1% sodium deoxycholate, 1 mM Na3VO4, 1 μg/ml leupeptin, 2 mM phenylmethanesulfonyl fluoride, 1 μg/ml aprotinin, 1 μg/ml pepstatin). Then, lysed cells were scraped off and kept at 4° for 15 min with gentle agitation. Lysis was followed by centrifugation of the lysates at 14000 g at 4° for 15 min to separate insoluble particles and debris. Subsequently, the lysate was incubated with 100 μl protein A/G beads (Biolinkedin, Shanghai, China) at 4° for 15 min to eliminate non-specific interactions. Then, 500 μl of lysate was incubated with 5 μg of antibodies and rotated at 4° overnight. After washing and elution, precipitated proteins were measured by western blot analysis.

### Statistical method

SPSS 15.0 software (SPSS, Inc., Chicago, IL, USA) was used to analyse the data, which are presented as the means ± SEM. Data were analysed with t-tests and one-way analysis of variance. Statistical significance was defined as p < 0.05.

## Results

### Histological changes in the airway of allergic asthma rats

The numbers of mucosal and submucosal inflammatory cells in lung tissue stimulated by ovalbumin were significantly augmented compared to the control group. Meanwhile, the airway wall and airway smooth muscle layer in asthma rats were markedly thicker than in the control group (Fig. 1A). Smooth muscle α-actin (α-SMA) immunofluorescence was performed to identify ASMCs in asthmatic rats. These results in the allergic asthma group are shown in Figure 1B. We observed that subcultured cells presented as primarily as fusiform and sometimes as polygonal arranged neatly with round nuclei located in the centre, and ASMCs was α-SMA positive.

**Figure. 1.**
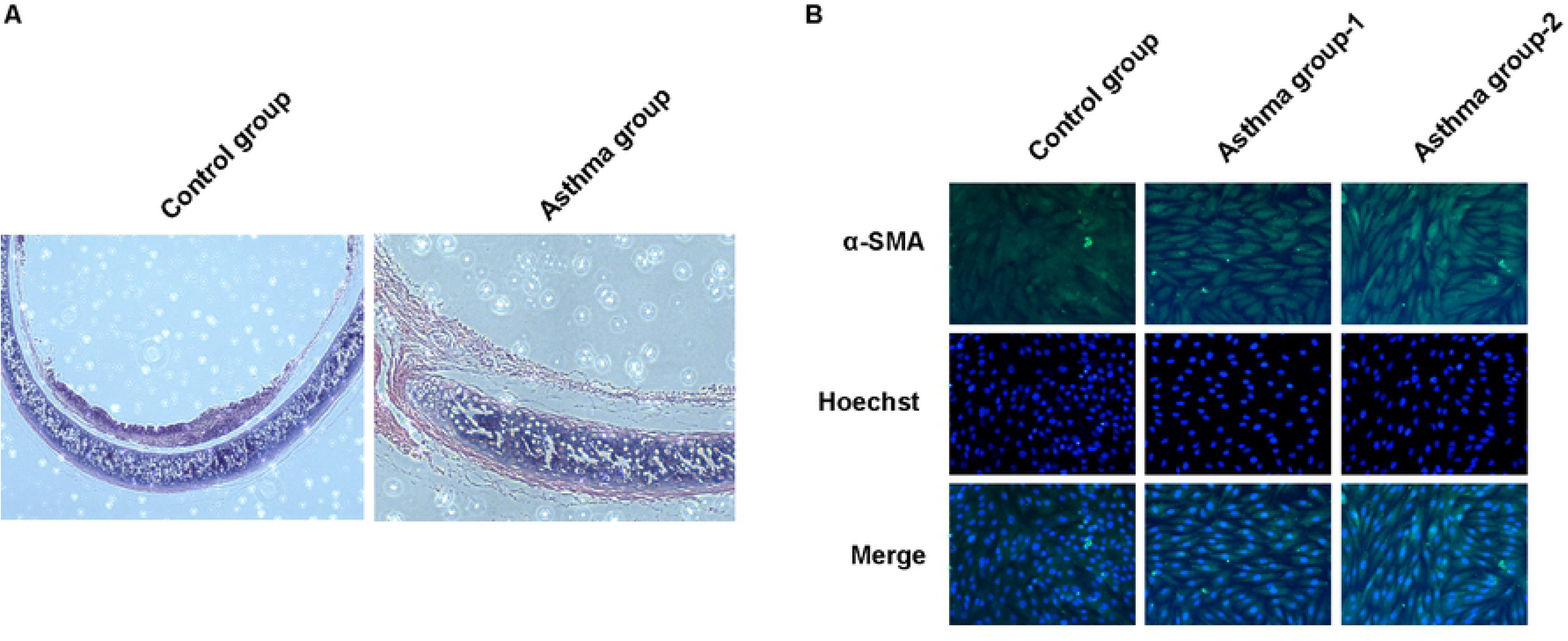
Histological changes in the airway of allergic asthma rats. (A) HE staining of lung tissue and bronchial wall in rats, ×40. The picture shows the lung tissue and airway wall constructions in control group (Left) and asthma group (Rigtht). (B) Representative immunofluorescent stained cells from trachea of wistar rats in the control group and asthma group. Upper: wistar rats ASM cells under the luminescence microscope following immunofluorescent identification by α-SMA staining. Magnification×400.

### Growth, migration and intracellular Ca2+ levels of ASMCs were significantly increased in ASMCs from asthmatic rats

As shown in Figure 2A and 2B, the viability and migration ability of ASMCs isolated from asthmatic rats were higher than ASMCs isolated from the control group. Meanwhile, ASMCs isolated from asthmatic rats displayed markedly increased Ca2+ concentrations compared to control cells (Fig. 3C).

**Figure. 2.**
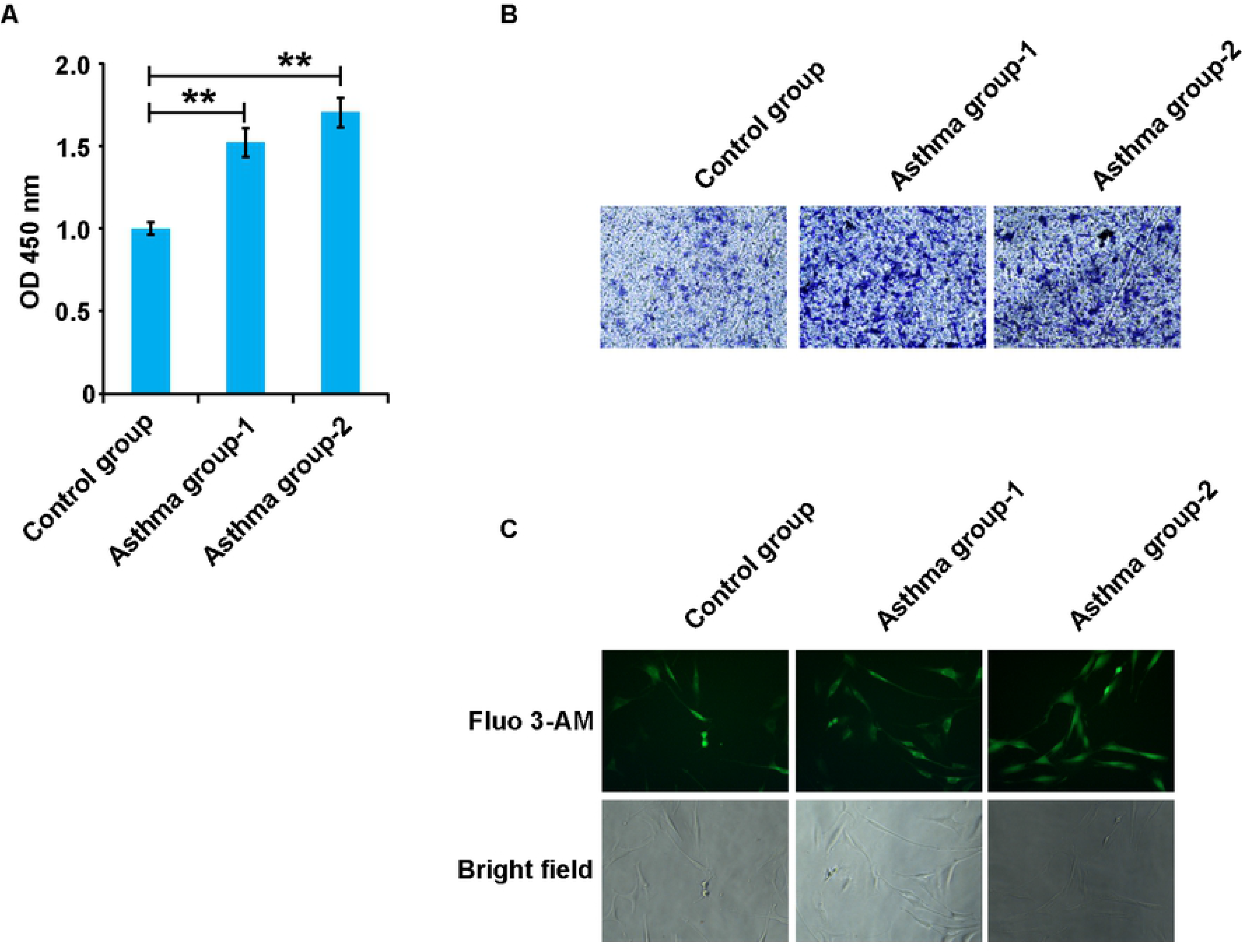
Growth, migration and intracellular Ca2+ levels of ASMCs were significantly increased in ASMCs from asthmatic rats. (A) The viability of ASMCs were detected after treatment with OVA(n=10). (B) The migration of ASMCs were detected after treatment with OVA(n=10). (C) The concentration of intracellular calcium in both asthma group and control group. Data are expressed as the mean±standard deviation. There was a significant difference between the two groups. **P<0.01 versus the control group.

**Figure. 3.**
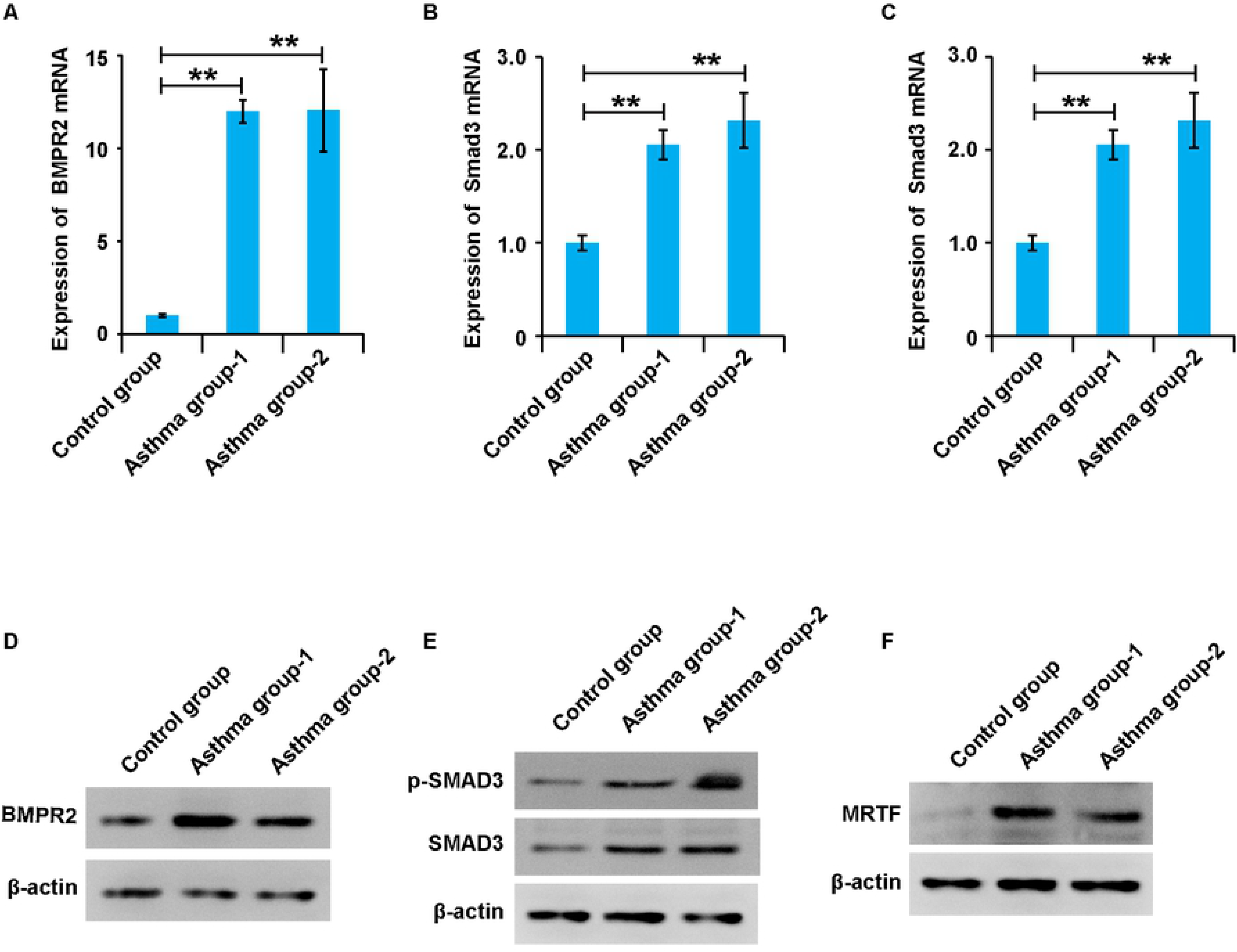
BMPR-II, SMAD3, and MRTF are upregulated in ASMCs from asthmatic rats. (A-C) The mRNA levels of BMPR-II, SMAD3, and MRTF in ASMCs was detected by qPCR in asthma group and control group. (D-E) The protein levels of BMPR-II, SMAD3, and MRTF in ASMCs was detected by Western blot in asthma group and control group. β-Actin was used as internal control. Data are expressed as the mean±standard deviation. **P < 0.01 was considered to indicate a statistically difference compared with the control.

### BMPR-II, SMAD3, and MRTF are upregulated in ASMCs from asthmatic rats

To explore the role of BMPR-II, SMAD3, and MRTF in asthma, we detected expression of these three factors in ASMCs from non-asthma and asthma model rats. qPCR results demonstrated that mRNA expression levels of BMPR-II, SMAD3, and MRTF were significantly upregulated in ASMCs from asthma model rats compared to non-asthma rats (Fig. 4A-C). Similar to protein expression patterns, western blots results showed that the protein expression levels of BMPR-II, SMAD3, and MRTF were significantly up-regulated in ASMCs from asthmatic rats (Fig. 4D-F). These results imply that high expression of BMPR-II, SMAD3, and MRTF might be associated with the progression of asthma. Expression of these three factors is known to be related to the pathogenesis of asthma, but whether there is an interaction between the three and joint regulation of ASMCs leading to airway smooth muscle remodelling occurs need further research.

**Figure. 4.**
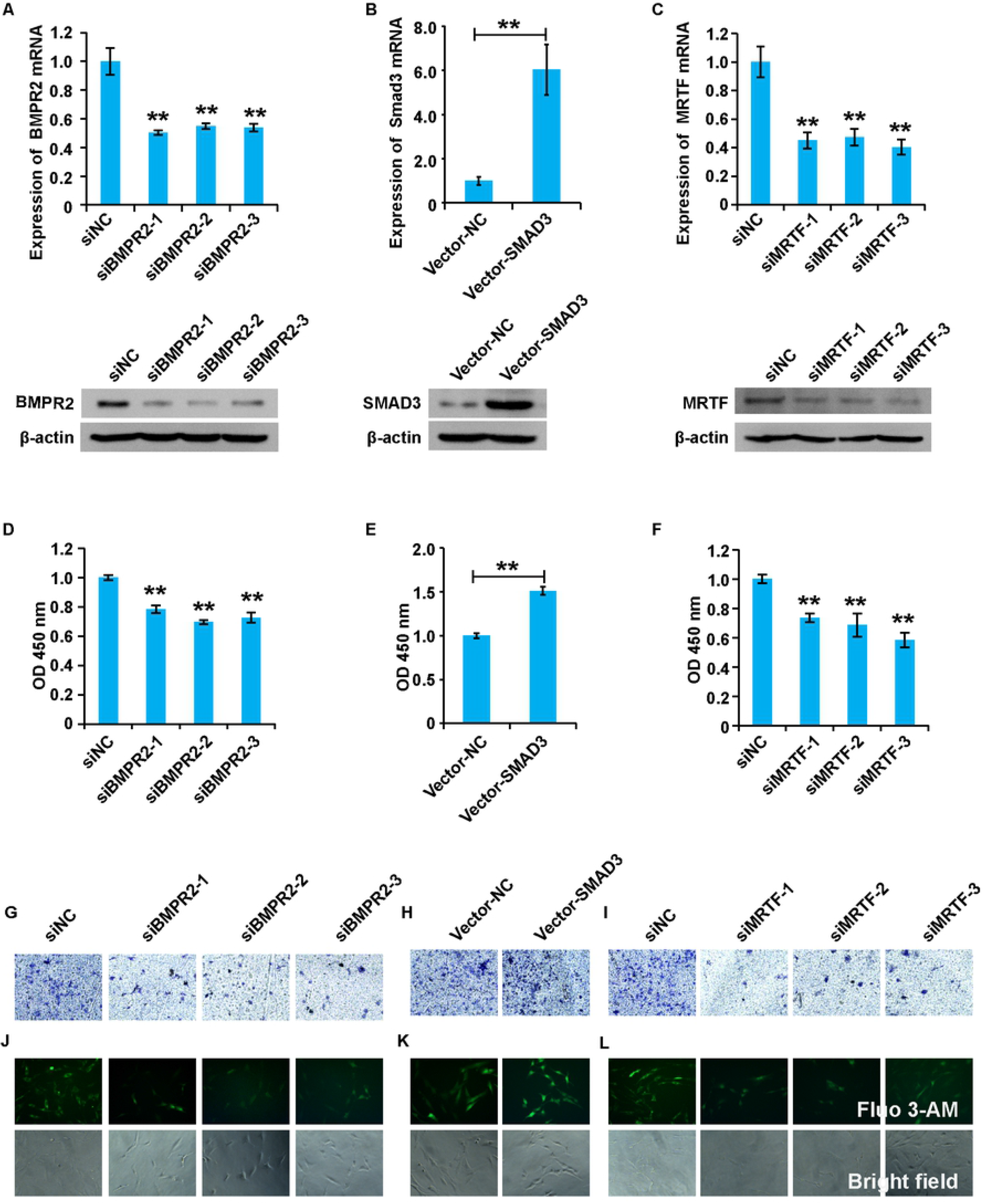
The BMPR-II, SMAD3 and MRTF complex affect cell viability, migration capacity, and intracellular Ca2+ concentration in asthmatic rats ASMCs. (A-C) qPCR and western blot assay analysis of the validity of siBMPR-II, vector-SMAD3 and siMRTF respectively. (D-F) CCK9 assay analysis of the viability of ASMCs with different treatments. (G-I) Transwell assay analysis of the migration of ASMCs with different treatments. (J-L) The concentrations of intracellular calcium in ASMCs with different treatments. β-Actin was used as internal control. Data are expressed as the mean±standard deviation. There was a significant difference between the two groups. **P<0.01 versus the control group.

### The BMPR-II, SMAD3 and MRTF complex affect cell viability, migration capacity, and intracellular Ca2+ concentration in asthmatic rats ASMCs

To further examine the effects of BMPR-II, SMAD3 and MRTF on the bioactivity of asthmatic ASMCs, siBMPR-II, pcDNA-SMAD3, siMRTF were designed and transfected into ASMCs, respectively. Overexpression and knockdown efficiency were validated by qPCR and western blot. First, qPCR and western blot analysis displayed that all three siBMPR-IIs efficiently inhibited BMPR-II expression in asthmatic ASMCs, and siBMPR-II-2 was chosen for subsequent experiments (Fig. 4A). CCK8, migration and Fluo-4AM analysis demonstrated that silencing siBMPR-II reduced cell viability, inhibited cell migration and attenuated intracellular Ca2+ concentration in ASMCs (Fig. 4D, G and J). Second, qPCR and western blot analysis revealed that pcDNA-SMAD3 significantly elevated SMAD3 mRNA and protein levels, respectively, in asthmatic ASMCs (Fig. 4B). CCK8, migration and Fluo-4AM analysis revealed that up-regulation of SMAD3 enhanced cell viability, promoted cell migration and elevated intracellular Ca2+ concentration in ASMCs (Fig. 4E, H and K). Third, qPCR and western blot analysis illustrated that all three siMRTFs efficiently inhibited MRTF expression in asthmatic ASMCs, and siMRTF-1 was chosen for use in subsequent experiments (Fig. 4C). CCK8, migration and Fluo-4AM analysis revealed that knockdown of siMRTF reduced cell viability, inhibited cell migration and decreased intracellular Ca2+ concentration in ASMCs (Fig. 4F, I and L). Collectively, these results demonstrated that BMPR-II, SMAD3 and MRTF affect cell viability, migration capacity, and intracellular Ca2+ concentration in asthmatic rat ASMCs.

### BMPR-II, SMAD3 and MRTF form a functional complex

Previous studies have shown there is a direct link among BMPR-II, SMAD3 and MRTF in asthma. Thus, we examined whether BMPR-II, SMAD3 and MRTF form a complex that regulates the bioactivity of ASMCs. To answer this question, we performed co-immunoprecipitation analysis, and results demonstrated that the anti-BMPR-II antibody could pull down SMAD3 and MRTF, uncovering cross talk among BMPR-II, SMAD3 and MRTF (Fig. 5). Taken together, these results demonstrate that BMPR-II, SMAD3 and MRTF form a complex.

**Figure. 5.**
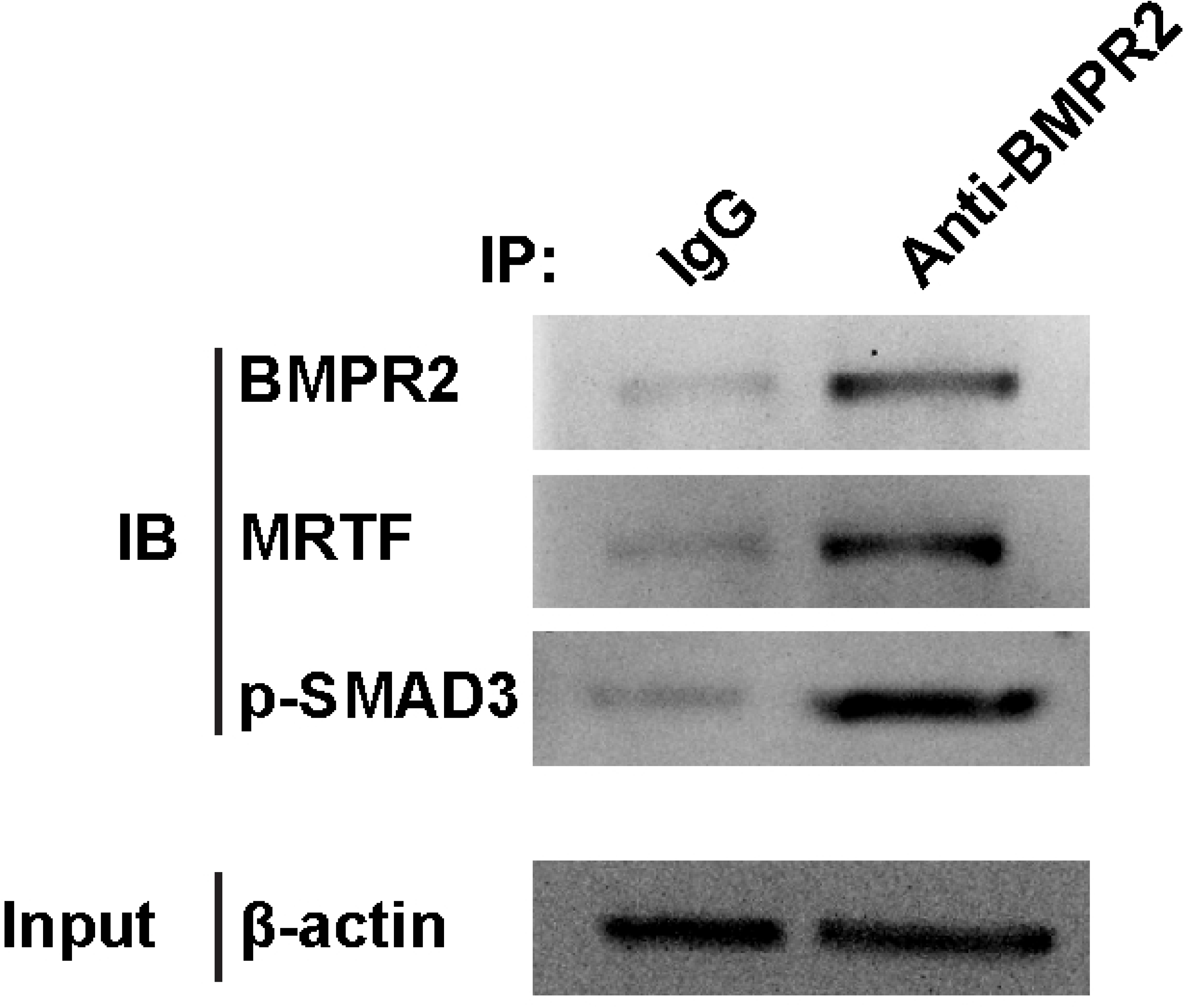
Interaction of BMPR-II, SMAD3 and MRTF. The pulling antibody and the blotting antibody indicate that SMAD3 could be pulled down by an anti-BMPR-II antibody. Meanwhile BMPR-II and SMAD3 both could be pulled down by an anti-MRTF antibody. Control immunoprecipitation was performed using pre-immune IgG. IP represents immunoprecipitation and IB indicates immunoblot.

### Effects of the BMPR-II-SMAD3/MRTF complex on proliferation and migration capacity

To confirm the physical or functional interaction among BMPR-II, SMAD3, and MRTF, we subsequently performed the following experiments. First, we divided ASMCs into four groups: siNC+pcDNA-NC, siBMPR-II+pcDNA-NC, siBMPR-II+pcDNA-SMAD3 and siBMPR-II+pcDNA-SMAD3+siMRTF groups. CCK8, migration and Fluo-4AM analysis revealed that any disruption to the BMPR-II-SMAD3/MRTF complex affected cell viability, migration and intracellular Ca2+ concentration in ASMCs (Fig. 6A-C). Collectively, these results demonstrate that the BMPR-II-SMAD3/MRTF complex affects cell viability, migration capacity, and intracellular Ca2+ concentration in asthmatic rat ASMCs.

**Figure. 6.**
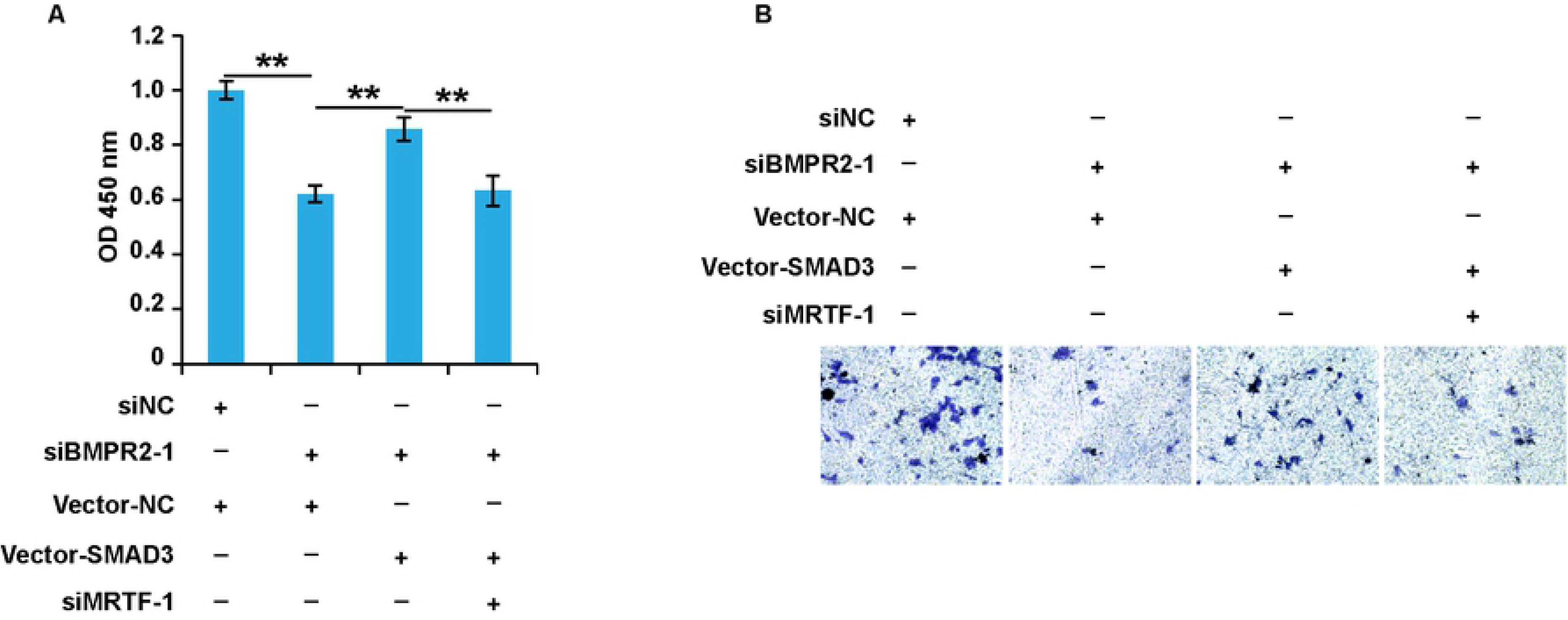
The effect of BMPR-II-SMAD3/MRTF signaling on the viability and migration of ASMCs. The viability of ASMCs with different treatment was detected. (B) The migration of ASMCs with different treatment was detected. Data are expressed as the mean±standard deviation(n=10). **P<0.01 was considered to indicate a statistically difference compared with the control.

## Discussion

Allergic asthma, an inflammatory respiratory disease, is defined as the inevitable result of chronic airway inflammation, airway hyperresponsiveness, and airway remodelling [18, 19]. Airway remodelling predominantly refers to changes in the airway wall components, which include epithelial shedding, fibroblast accumulation, peribronchial interstitial tissue changes, proliferation of the bronchovascular system, and increased smooth muscle mass [20]. These changes lead to airway stenosis and irreversible airway obstruction, which is the pivotal feature of recurrent asthma attacks [21]. ASM is the major tissue regulating airway lumen and predominantly mediating airway stenosis through contracting of airway smooth muscle, which were a critical target of historic asthma treatments [22]. Airway hyperresponsiveness is characteristic of strong contraction of the ASMCs when stimulated by a bronchoconstrictor [4]. However, not all patients with asthma exhibit an increase in ASM, and the pathological mechanism of ASM mediating the airway diameter remains unclear [23]. In recent years, increasing studies have shown that ASM remodelling occurs before airway inflammation, and ASMCs may represent the primary contributor to airway inflammation [24]. Airway inflammation promotes ASMC contraction and proliferation, whereas ASM remodelling aggravates airway inflammation by secreting a series of cytokines, forming a vicious cycle [25]. Current therapeutic drugs comprise corticosteroids and bronchodilators [26]. However, there are still patients who experience severe refractory asthma and develop resistance to these therapies [27]. Hence, it is imperative for us to identify new targets and to develop novel pharmacological interventions.

Recent studies have found that BMPR-II, SMAD3, and MRTF are all expressed in ASMCs in asthma models [14, 15, 28]. The contractility of ASMCs is largely dependent on intracellular Ca2+ concentrations; in addition, intrinsic increases in Ca2+ levels contribute to ASMC proliferation to some extent [29]. Moreover, prior studies demonstrated that instant Ca2+ transiently plays an important role in regulating cellular processes, such as proliferation and signal transduction [30]. Therefore, it is crucial to explore mechanisms for balancing the releasing of intracellular Ca2+ storage and extracellular Ca2+ entry to maintain Ca2+ homeostasis. Our experiments first revealed that BMPR-II, SMAD3, and MRTF are highly expressed in ASMCs in our asthma model. Furthermore, we found that inhibition of BMPR-II, SMAD3, and MRTF using siRNAs decreased proliferation and migration of ASMCs. Next, we confirmed that BMPR-II, SMAD3, and MRTF form a complex in our atopic asthma model, as evidenced by immunoprecipitation assays. All three molecules in this complex affect intracellular Ca2+ levels. Together with previous studies, these results led us to hypothesize that this complex may play a crucial role in the regulation of cellular bioactivities, and the interaction between these three molecules may modulate contraction and proliferation of ASMCs through inducing the release of Ca2+. Furthermore, all three factors were indispensable to the function of the complex.

To explore this interaction and specific functional mechanisms involved, we performed several additional experiments. First, we knocked down BMPR-II in asthmatic ASMCs and observed decreased proliferation and migration rates of ASMCs. Since all three factors were highly expressed in ASMCs and BMPR-II was inhibited by siBMPR-II, we subsequently selected pcDNA SMAD3 to overexpress SMAD3 to detect the interaction between SMAD3 and BMPR-II. As expected, we found that ASMCs treated with pcDNA SMAD3 exhibited increased proliferation and migration compared to the cells with BMPR-II inhibition. When cells were treated with siMRTF, cell viability and migration decreased compared to cells in the siBMPR-II-pcDNA SMAD3 group. However, there was no significant difference compared to ASMCs in the siBMPR-II group. Taken together, these findings support a physical and functional interaction among BMPR-II, SMAD3, and MRTF, which may involve regulation of proliferating and migration in ASMCs through mediating intracellular Ca2+ concentrations. However, this specific mechanism needs to be further studied.

## Conclusions

Thus, this study displays the functional complex, BMPR-II-SMAD3/MRTF signalling pathway, and further, revealed their huge role in the process of ASM remodelling, including the cell growth, migration and intracellular Ca2+ concentration and thus, indicates a potential of the complex in construction of the novel treatment for asthma.

## Author contributions

WG and LZ designed research; WG, JX, JL and YX performed experiments; WG, JX and LZ analyzed data, interpreted results and figures; WG drafted manuscript; WG, JX, JL, YX and LZ approved final version of manuscript.

## Acknowledgement

This work was supported by the National Natural Science Foundation of China (grant numbers: U1604161). The funding agencies had no role in study design, collection and analyses of data, decision to publish, or preparation of the article.

## Reference

1 Boulet LP, O’Byrne PM. Asthma and exercise-induced bronchoconstriction in athletes. The New England journal of medicine 2015, 372: 641–648

2 Mims JW. Asthma: definitions and pathophysiology. International forum of allergy & rhinology 2015, 5 Suppl 1: S2–6

3 Fehrenbach H, Wagner C, Wegmann M. Airway remodeling in asthma: what really matters. Cell and tissue research 2017, 367: 551–569

4 Chen J, Miller M, Unno H, Rosenthal P, Sanderson MJ, Broide DH. Orosomucoid-like 3 (ORMDL3) upregulates airway smooth muscle proliferation, contraction, and Ca(2+) oscillations in asthma. The Journal of allergy and clinical immunology 2018, 142: 207–218 e206

5 Saglani S, Lloyd CM. Novel concepts in airway inflammation and remodelling in asthma. The European respiratory journal 2015, 46: 1796–1804

6 Boulet LP. Airway remodeling in asthma: update on mechanisms and therapeutic approaches. Current opinion in pulmonary medicine 2018, 24: 56–62

7 Khan MA. Inflammation signals airway smooth muscle cell proliferation in asthma pathogenesis. Multidisciplinary respiratory medicine 2013, 8: 11

8 Benayoun L, Druilhe A, Dombret MC, Aubier M, Pretolani M. Airway structural alterations selectively associated with severe asthma. American journal of respiratory and critical care medicine 2003, 167: 1360–1368

9 Pascual RM, Peters SP. Airway remodeling contributes to the progressive loss of lung function in asthma: an overview. The Journal of allergy and clinical immunology 2005, 116: 477–486;quiz 487

10 Chu S, Zhang X, Sun Y, Liang Y, Sun J, Lu M, Huang J, et al. Atrial natriuretic peptide inhibits epithelial-mesenchymal transition (EMT) of bronchial epithelial cells through cGMP/PKG signaling by targeting Smad3 in a murine model of allergic asthma. Experimental lung research 2019, 45: 245–254

11 Silvestri L, Nai A, Dulja A, Pagani A. Hepcidin and the BMP-SMAD pathway: An unexpected liaison. Vitamins and hormones 2019, 110: 71–99

12 Groneberg DA, Witt H, Adcock IM, Hansen G, Springer J. Smads as intracellular mediators of airway inflammation. Experimental lung research 2004, 30: 223–250

13 Miyazawa K, Miyazono K. Regulation of TGF-beta Family Signaling by Inhibitory Smads. Cold Spring Harbor perspectives in biology 2017, 9

14 Kariyawasam HH, Xanthou G, Barkans J, Aizen M, Kay AB, Robinson DS. Basal expression of bone morphogenetic protein receptor is reduced in mild asthma. American journal of respiratory and critical care medicine 2008, 177: 1074–1081

15 Kumawat K, Koopmans T, Menzen MH, Prins A, Smit M, Halayko AJ, Gosens R. Cooperative signaling by TGF-beta1 and WNT-11 drives sm-alpha-actin expression in smooth muscle via Rho kinase-actin-MRTF-A signaling. American journal of physiology Lung cellular and molecular physiology 2016, 311: L529–537

16 Taiyab A, Holms J, West-Mays JA. beta-Catenin/Smad3 Interaction Regulates Transforming Growth Factor-beta-Induced Epithelial to Mesenchymal Transition in the Lens. International journal of molecular sciences 2019, 20

17 Zabini D, Granton E, Hu Y, Miranda MZ, Weichelt U, Breuils Bonnet S, Bonnet S, et al. Loss of SMAD3 Promotes Vascular Remodeling in Pulmonary Arterial Hypertension via MRTF Disinhibition. American journal of respiratory and critical care medicine 2018, 197: 244–260

18 Khan DA. Allergic rhinitis and asthma: epidemiology and common pathophysiology. Allergy and asthma proceedings 2014, 35: 357–361

19 Zissler UM, Esser-von Bieren J, Jakwerth CA, Chaker AM, Schmidt-Weber CB. Current and future biomarkers in allergic asthma. Allergy 2016, 71: 475–494

20 Manuyakorn W, Howarth PH, Holgate ST. Airway remodelling in asthma and novel therapy. Asian Pacific journal of allergy and immunology 2013, 31: 3–10

21 Schatz M, Rosenwasser L. The allergic asthma phenotype. The journal of allergy and clinical immunology In practice 2014, 2: 645–648;quiz 649

22 Gazzola M, Mailhot-Larouche S, Beucher C, Bosse Y. The underlying physiological mechanisms whereby anticholinergics alleviate asthma. Canadian journal of physiology and pharmacology 2018, 96: 433–44

23 Nair P, Martin JG, Cockcroft DC, Dolovich M, Lemiere C, Boulet LP, O’Byrne PM. Airway Hyperresponsiveness in Asthma: Measurement and Clinical Relevance. The journal of allergy and clinical immunology In practice 2017, 5: 649–659 e642

24 Noble PB, Pascoe CD, Lan B, Ito S, Kistemaker LE, Tatler AL, Pera T, et al. Airway smooth muscle in asthma: linking contraction and mechanotransduction to disease pathogenesis and remodelling. Pulmonary pharmacology & therapeutics 2014, 29: 96–107

25 West AR, Syyong HT, Siddiqui S, Pascoe CD, Murphy TM, Maarsingh H, Deng L, et al. Airway contractility and remodeling: links to asthma symptoms. Pulmonary pharmacology & therapeutics 2013, 26: 3–12

26 Chung KF. Clinical management of severe therapy-resistant asthma. Expert review of respiratory medicine 2017, 11: 395–402

27 Castillo JR, Peters SP, Busse WW. Asthma Exacerbations: Pathogenesis, Prevention, and Treatment. The journal of allergy and clinical immunology In practice 2017, 5: 918–927

28 Wang M, Li H, Zhao Y, Lv C, Zhou G. Rhynchophylline attenuates allergic bronchial asthma by inhibiting transforming growth factor-beta1-mediated Smad and mitogen-activated protein kinase signaling transductions in vivo and in vitro. Experimental and therapeutic medicine 2019, 17: 251–259

29 Reyes-Garcia J, Flores-Soto E, Carbajal-Garcia A, Sommer B, Montano LM. Maintenance of intracellular Ca2+ basal concentration in airway smooth muscle (Review). International journal of molecular medicine 2018, 42: 2998–3008

30 Chen YF, Cao J, Zhong JN, Chen X, Cheng M, Yang J, Gao YD. Plasma membrane Ca2+-ATPase regulates Ca2+ signaling and the proliferation of airway smooth muscle cells. European journal of pharmacology 2014, 740: 733–741

